# Does Music Training Improve Inhibition Control in Children? A Systematic Review and Meta-Analysis

**DOI:** 10.1101/2023.02.08.527718

**Authors:** Kevin Jamey, Nicholas E. V. Foster, Krista L. Hyde, Simone Dalla Bella

**Affiliations:** International Laboratory for Brain, Music and Sound Research (BRAMS), University of Montreal Centre for Research on Brain, Language and Music (CRBLM), Montreal, Quebec, Canada

**Keywords:** inhibition control, music training, child development, meta-analysis, skill transfer

## Abstract

Inhibition control is an essential executive function during children’s development, underpinning self-regulation and the acquisition of social and language abilities. This executive function is intensely engaged in music training while learning an instrument, a complex multisensory task requiring monitoring motor performance and auditory stream prioritization. This novel meta-analysis examined music-based training on inhibition control in children. Records from 1980 to 2023 yielded 22 longitudinal studies with controls (N = 1734), including 8 RCTs and 14 others. A random-effects meta-analysis showed that music training improved inhibition control (medium ES) in the RCTs and the superset of twenty-two longitudinal studies (small-to-medium ES). Music training plays a privileged role compared to other activities (sports, visual arts, drama) in improving children’s executive functioning, with a particular effect on inhibition control. We recommend music training for complementing education and as a clinical tool focusing on inhibition control remediation (e.g., in autism and ADHD).

## Main

Everyday tasks like talking, walking, and playing rely heavily on integrating our senses and motor skills. Learning to play a musical instrument likewise depends on the effective coordination of physical movements and auditory signals to produce music. Mastery in musical training develops through repetitive practice, consistently honing the connection between perception, action, and cognition (Baer et al., 2013; Dalla Bella et al., 2024; Repp, 2010). Music training is a rich and highly engaging experience for children and can potentially strengthen cognitive skills essential for healthy child development. Merely listening to music elicits activation in the brain’s reward circuits in both children (Fasano et al., 2022) and adults (Mas-Herrero et al., 2021; Vuust et al., 2022; Zatorre, 2015). Being internally motivated during learning experiences increases learning capacity and efficiency (Ryan & Deci, 2017), and this greater engagement is reflected in increased electrical brain activity following musical training (Kraus et al., 2014). The external setting in which learning occurs also plays an important role. Training music in a social environment increases positive feelings of bonding through shared emotions and group synchrony (Nummenmaa et al., 2021). In addition to being pleasurable and social, music training also engages a broad set of skills including executive functioning (Degé & Frischen, 2022; Rodriguez-Gomez & Talero-Gutiérrez C., 2022), language/communication (Gordon et al., 2015; Jäncke, 2012; Patel, 2008; Poeppel & Bergelson, 2008), and socio-emotional functioning (Gaudette-Leblanc et al., 2021; Gerry et al., 2012). Given the wealth of different processes engaged in music training, music emerges as a precious activity for studying skill transfer effects. Some transfers are relatively close to the trained ability and considered near transfers, for example, between music training and sensorimotor integration. Conversely, some are considered more indirect or distantly related to the trained ability and are referred to as far transfers. Some examples of far transfer from music training include effects on executive functioning, speech, and quality of life.

Training of cognition through music has attracted considerable research attention over the last two decades and is currently part of an ongoing debate. Sala and Gobet (2020) conducted a multilevel meta-analysis on music training’s effects on non-verbal ability, verbal ability, memory, and executive function speed, reporting no transfer effect (g = 0.06). However, the study faced criticisms due to methodological concerns, including non-equidistant comparisons and the inclusion of studies lacking pre-training scores (Bigand & Tillmann, 2021). Bigand and Tillmann (2021) re-analyzed the data entered in Sala and Gobet’s meta-analysis, adjusting criteria to address these issues, finding a modest yet dependable effect of music training (g = 0.23). A subsequent meta-analysis by Román-Caballero et al. (2022) similarly indicated a positive impact of music training on cognitive and academic skills (g = 0.28).

In summary, current evidence suggests a small effect size for far transfer from music training to cognition. These results encourage future meta-analytic work on music training and far skill transfers to cognitive domains. Importantly, Bigand & Tillmann (2021) highlight that meta-analyses are a work in progress and require care to avoid unintended biases in selecting studies and conditions for analysis — especially in cognitive science, where experimental methodologies may vary significantly across studies. Although several meta-analyses have been published examining far transfer from music training to various cognitive abilities, no meta-analysis has narrowed the focus to a specific executive function. In 2022, two systematic reviews were carried out investigating the effects of music training on a variety of executive functions and found that inhibition control was the most predominantly studied and showed the most reliable impact (Degé & Frischen, 2022; Rodriguez-Gomez & Talero-Gutiérrez C., 2022). Our meta-analysis builds on previous work by targeting a specific executive function, inhibition control, which may serve as a key driver of skill transfer from music training to executive functioning. Compared to a more holistic approach, one advantage of this method is its theory-driven nature, which connects the specific cognitive function expected to be influenced by music training with the underlying mechanisms.

Meta-analyses present many advantages compared to individual studies, especially when studies involve low small samples of participants. They enhance scientific progress by systematically synthesizing knowledge from numerous studies and yield accurate effect size estimates by combining data from independent sources, boosting statistical power and generalizability. By aggregating a larger sample size, meta-analyses can unveil patterns, trends, and relationships that might remain hidden in individual studies due to their inherent limitations. Moreover, they pinpoint variations and biases in research, allowing exploration of factors impacting result consistency. By offering a comprehensive evidence overview, meta-analyses guide discussions, inform decisions, and direct future research, promoting objective and rigorous knowledge advancement.

In causal inference, the parallel-arm Randomized Control Trials (RCT) design with an active control group is a standard method (Kendall, 2003). Randomizing participant assignment mitigates selection biases, while an active control group clarifies effects in the treatment group. Careful control group design, ensuring similarity in participation without targeting the experimental function, is crucial. Poor design choices can equate to a passive group, highlighting the importance of active control groups for interpreting findings accurately. These groups control for placebo effects, expectancy effects, and non-specific changes, enhancing result interpretation (Foroughi et al., 2016; von Bastian & Oberauer, 2014).

Recent short longitudinal experiments using active control groups and an RCT approach with targeted outcomes in 4 to 8-year-old children show converging and promising evidence of far skill transfer effects from music training to inhibition control (Frischen et al., 2019; Patscheke et al., 2019; Williams & Berthelsen, 2019; 480 min-960 min training time over 12-16 weeks). These studies highlight a growing interest in theory-based and mechanism-driven RCTs with targeted outcomes and active controls in music training. The starting age for music training is essential for benefiting from sensitive learning periods during brain development. Studying children during a specific time window to develop their cognitive and musical abilities (4 to 7 years of age) may help increase the impact of shorter music training programs on brain plasticity (Bailey & Penhune, 2013; Best & Miller, 2010). Music training is a highly complex activity and studying it through the lens of specific mechanism-based transfer effects is beneficial for understanding and interpreting music-induced skill transfers.

Music training strongly engages executive functions, particularly inhibition control, which encompasses response inhibition—the ability to halt repetitive actions—and attentional inhibition, which involves filtering out irrelevant stimuli (Tiego et al., 2018). Inhibition control is the earliest and fastest-developing executive function, emerging around two years of life (Best & Miller, 2010; Senn et al., 2004; Wiebe et al., 2008). In early childhood, inhibition control shows a quick improvement curve that becomes more gradual later in childhood (Best & Miller, 2010; Carlson, 2014; Nesbitt et al., 2015). Inhibition control is important for academic achievement (Allan et al., 2014; Liu et al., 2015; Wilkinson et al., 2020), goal-directed behavior (Allom et al., 2016), and reducing compulsive behavior during childhood (Fogel et al., 2019). In the early phases of music training, children grasp starting/stopping actions and use independent limbs on instruments like the piano or xylophone. They manage perceptual-motor coordination while producing musing, all demanding inhibition control. This ability is vital for attention in ensemble performance, responding to cues, and coordinating rhythmic patterns among musicians (Keller et al., 2014). Music training demands inhibition control for self-monitoring (intonation correction, error detection) and adapting one’s performance to changes in the musical structure, both global (key, tempo) and local (dynamics, polyrhythms) during joint action (Jentzsch et al., 2014; Okada & Slevc, 2019; Vuust et al., 2006, 2011; Zuk et al., 2014). Empirical data strongly support inhibitory control’s role in music training. Multiple RCT studies, many of which are included in this meta-analysis, show enhanced cognitive skills in children: inhibition control compared to visual arts, Lego, motor, sports training; set shifting, short-term memory over after-school programs; planning over visual arts; working memory compared to free play (Bolduc et al., 2021; Bowmer et al., 2018; Bugos & DeMarie, 2017; Frischen et al., 2019; Guo et al., 2018; Holochwost et al., 2017; Jaschke et al., 2018; Moreno et al., 2011).

Interestingly, some studies show a relation between specific music components and a selected cognitive function, such as inhibition control. For example, Frischen et al. (2019) showed that a rhythm-based music program improved inhibition control, whereas a program focused on pitch-based components of music did not. The observed effect may be due to the cognitive demands required during motor synchronization to a beat. In a similar study, pitch-based music training showed a greater impact on phonological awareness than rhythmic-based music training (Patscheke et al., 2019). The authors attribute this effect to a relationship between the elementary units of music (musical notes) and those of language (phonemes), which is in line with Patel’s OPERA hypothesis (Patel, 2011). Similarly, there is evidence that the entrainment of a regular rhythm in a training protocol strengthens emotional regulation and cognitive flexibility in children via “neural entrainment” (Williams, 2018). These findings highlight the effectiveness of targeting specific outcomes based on theory and mechanisms for studying skill transfers in music training.

Two key theoretical frameworks have helped elucidate the potential mechanisms behind skill transfer from music training. First, the OPERA hypothesis proposed by Patel (2011) places attention at the center of the training process; attention would be improved through intense repeated practice, involving, for example, training melody and rhythm perception and production and paving the way to transfer to language-related functions. One of the mechanisms by which music training transfers to verbal ability may be through the mediating effect of attentional regulation, a sub-function of inhibition control (Tiego et al., 2018). Second, the Dynamic Attending Theory explains that processing a temporally regular stimulus, like the underlying beat of music, can be modeled by the “entrainment” of internal attentional oscillations (Jones, 1976; Large & Jones, 1999). More recent work has provided the neurophysiological foundations of this theory using oscillatory neural dynamics (Fujioka et al., 2012; Nijhuis et al., 2021; Nozaradan, 2014, 2019). In this model, music with a regular beat provides a highly coherent structure, making it possible to form predictions about future events further than everyday temporal processes like speech (Dalla Bella et al., 2013). Training rhythmic abilities through music may optimize the “neural entrainment” of computations recruited by other music-related operations, like executive functioning or speech and socioemotional processing. These findings show that considering the roles of specific music components and the mechanisms by which they are implemented in the brain can provide significant insights for understanding skill transfer effects from music training.

This meta-analysis study sought to assess the effect of musical training on inhibition control in a developmental population for the first time. To this aim, we reviewed existing studies in children during a critical developmental age range for transfer effects from music training to inhibition control using a meta-analysis technique. To our knowledge, no meta-analysis on music training and inhibition control has been conducted for longitudinal research designs. Our meta-analysis is particularly valuable as it calculates effect size in RCT studies having active control groups. To assess the generalizability of these findings, we also examine all existing parallel-arm longitudinal work, including quasi-experimental designs.

The main objective of our study is to assess the existing causal evidence that music training administered in children leads to skill transfer to a non-musical ability and, more specifically, to improve inhibition control. The secondary objective is to examine whether this causal evidence is generalizable across real-world training contexts. The third objective is to investigate if the expected skill transfer from music training to inhibition control depends on age, training intensity, training context, and randomization procedure. Our specific aims and hypotheses are described as follows:

*Aim 1:* To determine if music training increases inhibition control in children by assessing RCT designs with active control training conditions.

*Aim 2:* To determine if music training increases inhibition control in children in an ecologically valid sample of longitudinal music training programs, including quasi-experimental designs.

Based on the outcomes of systematic reviews on executive functions and music training (Degé & Frischen, 2022; Rodriguez-Gomez & Talero-Gutiérrez C., 2022), we expected to find an increase in inhibition control because of participating in a music training program for *Aim 1* and *Aim 2*.

*Aim 3:* To examine if training inhibition control through music training is dependent on the following moderating variables:

a. Age. Based on work by Penhune (2011) on critical learning periods, we expected that children in this meta-analysis would benefit equally from music training because the developmental window (4-8 years of age) is in the range of the critical brain plasticity window for music.
b. Training intensity. Based on the prediction that repeated music practice generates greater skill transfer (Patel, 2011), we expected the programs with the most prolonged training time to produce the most significant increases in inhibition control.
c. Training context. Since more formal and individualized training makes children more accountable for their progress, we expected programs with more individualized training components and specialized institutions to be more effective than school-based programs.
d. Randomization. We expected music training would exhibit greater effects in non-randomized studies due to the pre-existing social cohesion of school classes.

## Results

Twenty-two studies (1734 participants across studies) were included in this review and meta-analysis. Sample sizes were calculated based on the data analyzed during pre-and post-training testing sessions. All studies reported data of equal sample sizes at the pre-training and post-training time points. The mean age across studies included in the meta-analysis was five years and eight months, ranging between three and eleven years. All studies included boys and girls, with a reported average of 54.3% girls, ranging between 40.9% to 71.8% across all studies. Eight studies used randomized group assignment at the participant level and had active control training conditions (Bolduc et al., 2021; Bowmer et al., 2018; Bugos & DeMarie, 2017; Degé et al., 2020; D’Souza & Wiseheart, 2018; Frischen et al., 2019, 2021; Moreno et al., 2011). Some studies were conducted in a school context with existing classroom groups, which did not permit participant-level randomization (see Supplemental Table F for study-level detail about randomization procedures; for the complete list of control group conditions, see Supplemental Table C). Eight studies were considered randomized control trial designs, representing 36% of the studies in this meta-analysis. Calculation of per-group attrition between commencement of training and completion of post-training measures was possible for 90% of the studies. A mixed-effects analysis of these attrition values showed no difference between the music training and control groups across studies (p = .93). Individual effect sizes and variances were calculated for each study’s most suitable inhibition control measure (see Table 1) and are shown in the forest plot of Figure 1.

**Figure 1.**
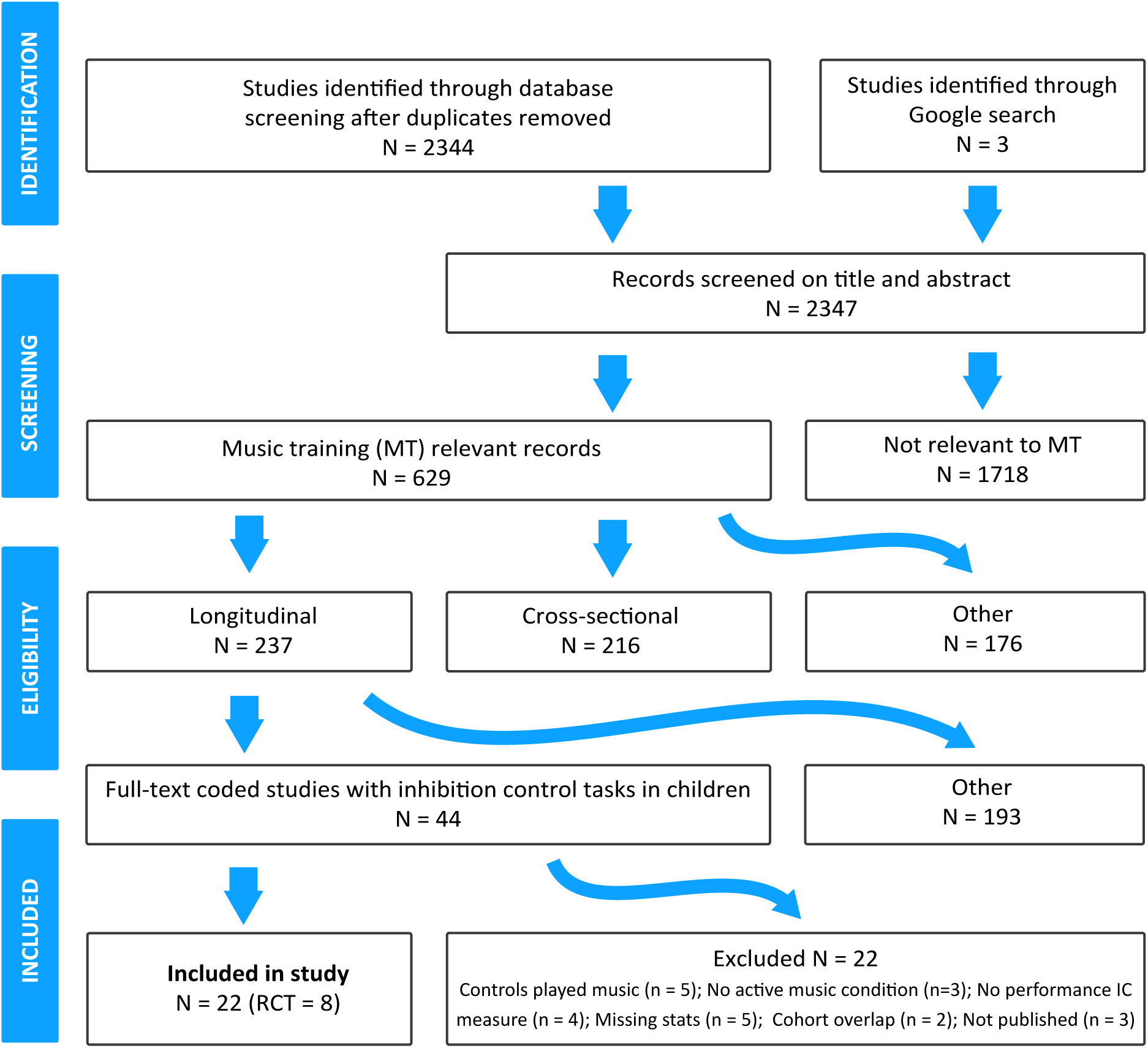
PRISMA Chart Showing the Number of Studies Included in the Meta-Analysis

**Table 1.** Results of Moderators of Interest.

Results of Moderators of Interest
| Moderator | RCTs | RCTs and Quasi-Experimental |
| --- | --- | --- |

|  | g | SE | CI | p-value | g | SE | CI | p-value |
| --- | --- | --- | --- | --- | --- | --- | --- | --- |
| Age | - | 0.13 | -0.32 – | .68 | 0.07 | 0.06 | -0.05 – 0.19 | .22 |
|  | 0.06 |  | 0.20 |  |  |  |  |  |
| Training | 0.05 | 0.13 | -0.20 – | .69 | - | 0.07 | -0.15 – 0.11 | .74 |
| Intensity |  |  | 0.30 |  | 0.02 |  |  |  |
| Intervention | 0.12 | 0.11 | -0.09 – | .24 | - | 0.07 | -0.26 – 0.00 | **.047\*** |
| Setting |  |  | 0.33 |  | 0.13 |  |  |  |
| Randomization |  |  |  |  | 0.06 | 0.07 | -0.08 – 0.19 | .41 |
| Procedure |  |  |  |  |  |  |  |  |

**Table 2.**
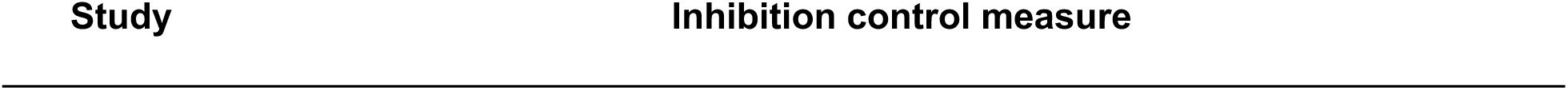
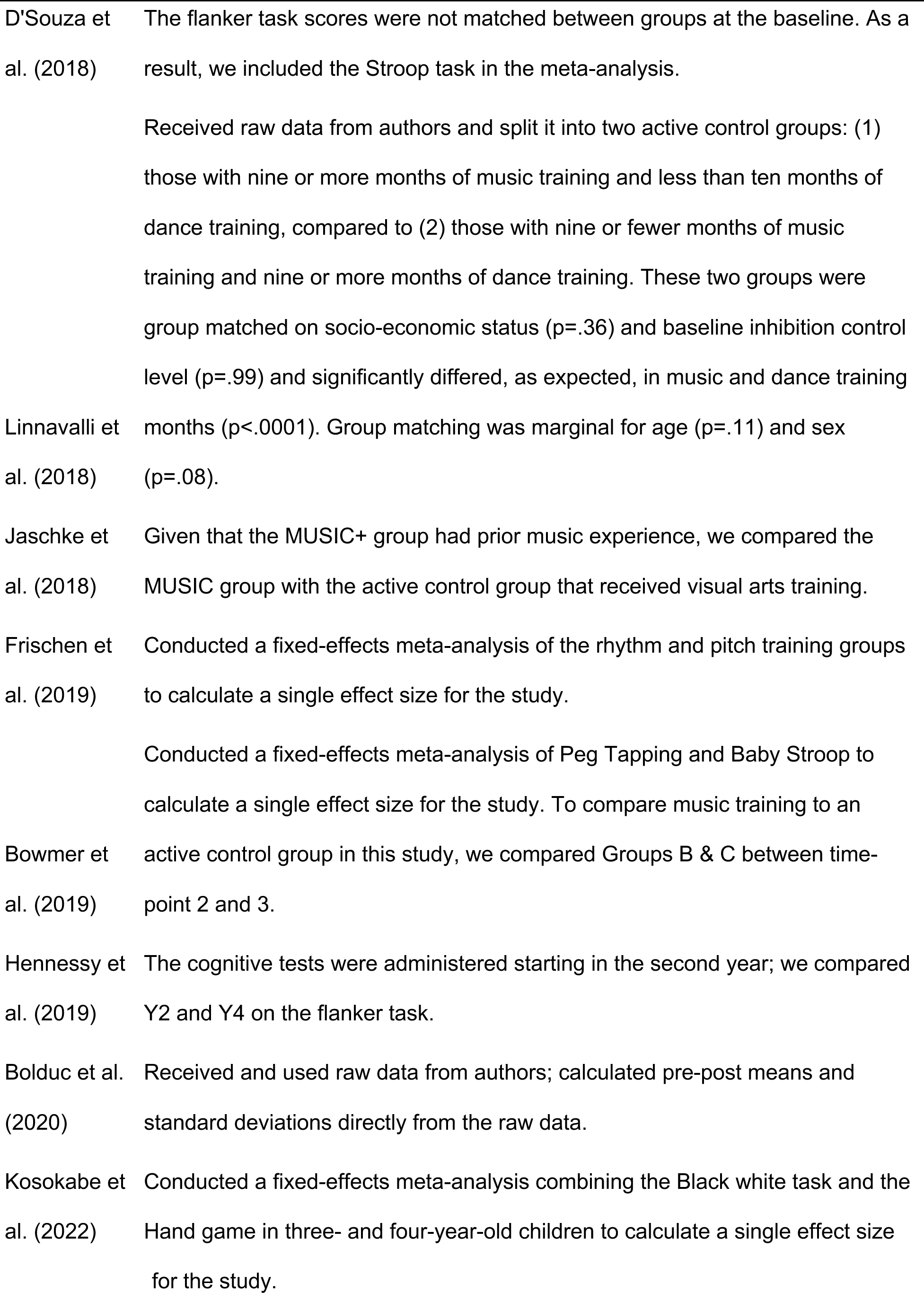
Decisions About Inhibition Control Measures Included in the Meta-Analysis.

### The effect of randomized control trials (active control) of music training on inhibition control

A random effects meta-analysis of 8 RCT studies indicated a significant overall effect size of g = 0.60 (SE = 0.11; CI = 0.39-0.82; p < .0001). This moderate-to-large effect indicates that inhibition control improves in children who participated in RCTs of music training compared to non-musical active control training. Individual effect sizes for studies in this analysis are shown in Figure 2. Results were robust when the value of rho was set to .001, yielding g = 0.60 (SE = 0.15; CI = 0.31-0.90; p < .0001) or rho = .999, yielding g = 0.55 (SE = 0.11; CI = 0.33-0.77; p < .0001).

**Figure 2.**
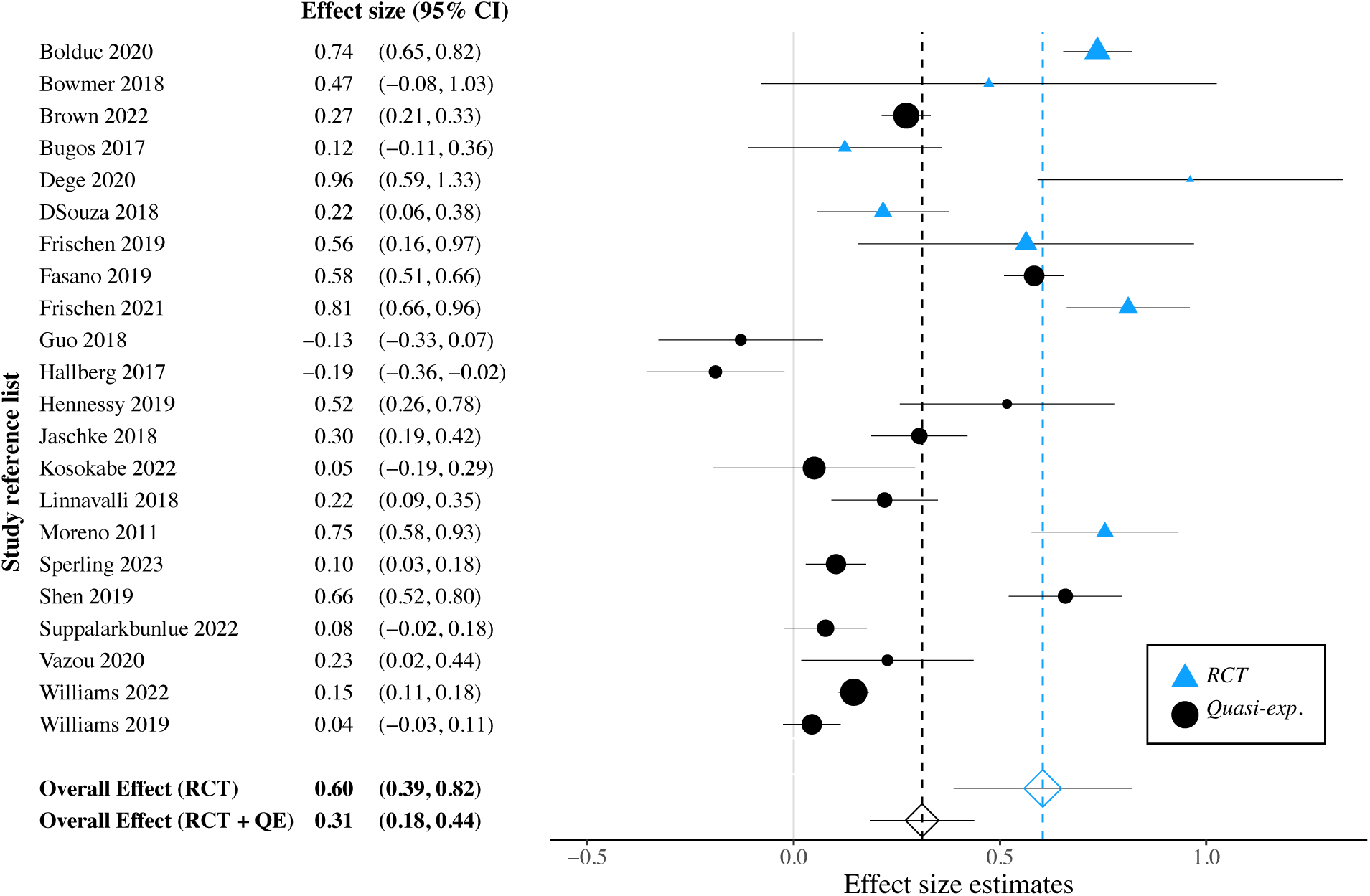
Forest plot of effect sizes (Hedges’ g) showing that music training improves inhibition control Note: Positive values to the right of the dotted line at 0 on the X-axis indicate better inhibition control after training in the music group

### The effect of longitudinal music training on inhibition control (active and passive control)

A random effects meta-analysis including all 22 longitudinal studies indicated a significant overall effect size of g = 0.31 (SE = 0.07; CI = 0.19-0.44; p < .0001). This small-to-moderate effect indicates that inhibition control improves in children who participated in music training programs compared to non-musical conditions (active and passive combined). Individual effect sizes for studies in this analysis are shown in Figure 2. All studies except Guo et al. (2018) and Hallberg et al. (2017) had a positive effect size. Results were robust when the value of rho was = .001, yielding g = 0.30 (SE = 0.07; CI = 0.16-0.45; p < .0001) or rho = .999, yielding g = 0.32 (SE = 0.07; CI = 0.19-0.45; p < .0001).

### Results of moderators of interest

Individual random effects meta-analyses were conducted in the set of 8 RCTs and the 22 longitudinal studies for age, training intensity (total minutes), intervention setting (in a school environment or not), and randomization (participant level or not). A moderating effect of the intervention setting was statistically significant (p = .047). All other analyses showed non-significant effects for these moderators of interest (p ≥ .22; see Table 1 for analysis statistics).

### Measuring between-study heterogeneity

Heterogeneity between studies was anticipated in this analysis due to variations in inhibition control measures available in each study, the characteristics of musical training and control groups, and differences in sample demographics such as age. The I^2^ estimate of inter-study heterogeneity was 0.0% for the analysis of RCT studies and 27.44% for all longitudinal studies. These values indicate a medium to low degree of variability between the studies (J. P. T. Higgins et al., 2019). Given that they have more homogeneous methodologies, lower variability for RCTs is expected.

### Small study bias

The potential for a small study bias was assessed by visually examining the symmetry of a funnel plot of the superset of longitudinal parallel-arm studies (Figure 3). No asymmetry in the funnel plot was observed, consistent with the absence of small study bias. Subsequently, Egger’s regression test confirmed a lack of asymmetry for the RCTs analysis and the superset of longitudinal parallel-arm studies (p > .33).

**Figure 3.**
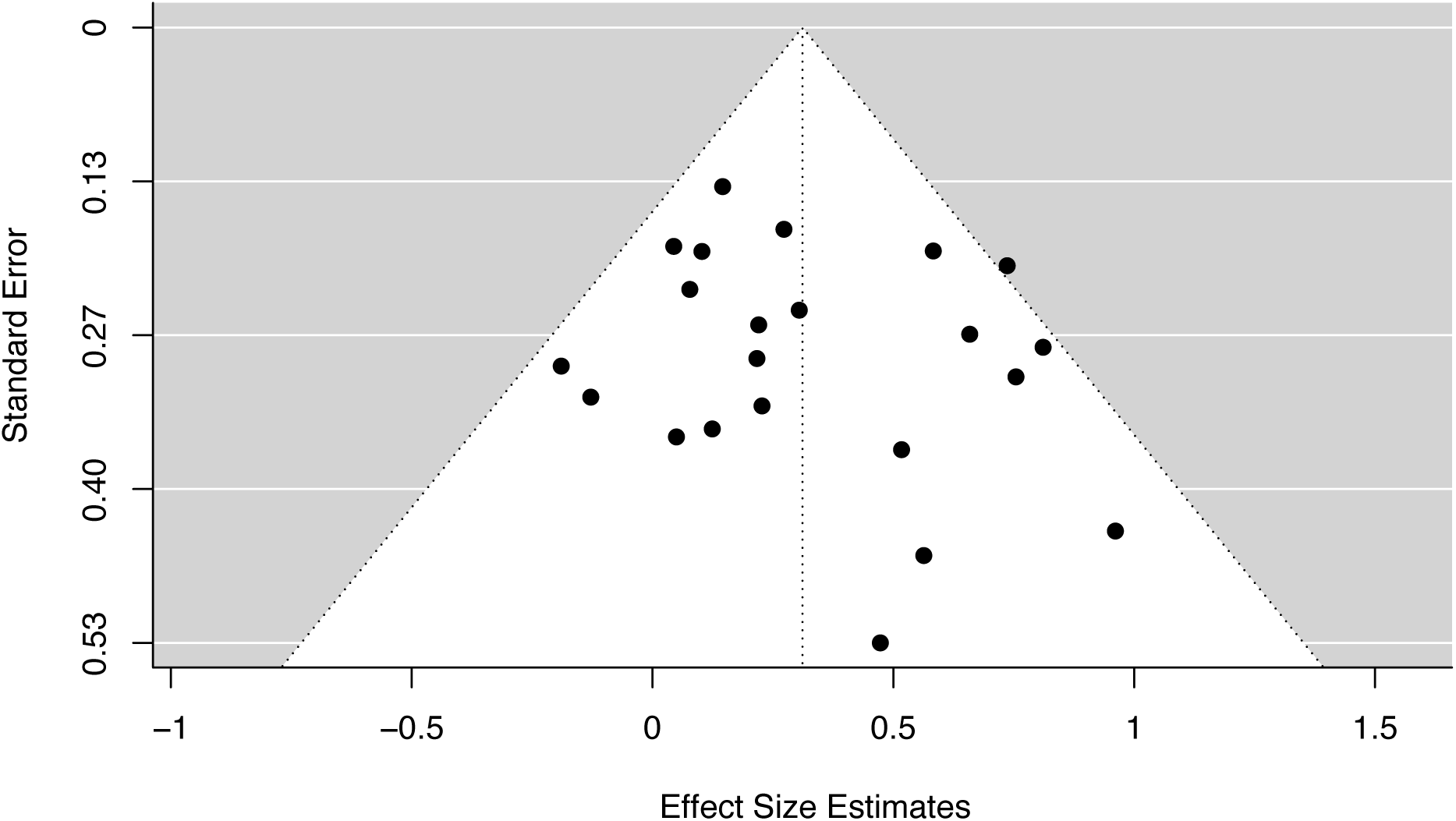
Funnel plot showing no small study bias when comparing effect sizes with standard errors for the superset of longitudinal parallel-arm studies Note: The dotted line represents a 95% CI

## Discussion

This meta-analysis aimed to assess for the first time the effect of music training in children on inhibition control, an essential executive function for child development (Allan et al., 2014; Liu et al., 2015; Wilkinson et al., 2020). This meta-analysis included studies between January 1980 and December 2023 in 3 to 11-year-old children. We show converging evidence that music training improves inhibition control with a moderate-to-large effect size estimate across 8 RCTs (g = 0.60) and a small-to-moderate effect size estimate in a superset of 22 longitudinal studies (g = 0.31), including the 8 RCTs. These results confirm skill transfer from music training to inhibition control, an effect hinted at by previous systematic reviews (Degé & Frischen, 2022; Rodriguez-Gomez & Talero-Gutiérrez C., 2022). Compared to most intervention effect sizes in education, this meta-analytic work’s RCT effect size estimate is particularly interesting, according to Hattie’s (2008) barometer of influence. After analyzing over 800 meta-analyses, Hattie (2008) reported that the median value of intervention effect sizes in education is 0.40. The author considers any effect size above that threshold to have a practical educational use. These findings also indicate that music training is more impactful in improving inhibition control than other types of activity aiming to generate skill transfer to inhibition control, such as video games. Effect size estimates for video games range between g = 0.27 (Sala et al., 2018) and g = 0.31 (Bediou et al., 2018). Our RCT findings suggest music training presents a unique advantage in training inhibition control compared to any other training program. Our findings highlight that larger effect sizes can be identified when the analytic criteria focus on a specific theory-driven research question.

Inhibition control is an important cognitive function in learning and playing a musical instrument. Music training requires inhibition control for coordinating sensorimotor actions in time with music and when playing in an ensemble with other musicians (Jentzsch et al., 2014; Okada & Slevc, 2019; Vuust et al., 2006, 2011; Zuk et al., 2014). Music training encompasses mechanisms for perceiving core musical elements (pitch, rhythm, timbre), motor control, auditory-motor integration, reward, and social interaction. It can be studied holistically or by isolating driving mechanisms (Frischen et al., 2019; Patscheke et al., 2019). Holistic training addresses the effect of training all music components, fostering connections throughout the brain, and broadly generalizing skills. Alternatively, specific musical elements like rhythm or pitch can be targeted to explore causality in transferring to distinct skills such as inhibition control or phonological awareness. The present analysis differs from previous meta-analyses because it concentrates on a single executive function, inhibition control, potentially enhancing sensitivity, and elucidating transfer implications and mechanisms.

Our findings add to the ongoing debate about far skill transfer effects in music training, providing a larger effect size estimate than previous meta-analyses involving executive functioning (Bigand & Tillmann, 2021; Román-Caballero et al., 2022; Sala & Gobet, 2020). Delineating near and far skill transfer in music training is challenging (Barnett & Ceci, 2002). Some view enhancements in non-musical executive functions as far transfer (D’Souza & Wiseheart, 2018; Miendlarzewska & Trost, 2013), while others perceive it as skill transfer within similar domains (Diamond & Lee, 2011; Frischen et al., 2021). Music engagement employs multiple executive functions (Jäncke, 2009; Okada & Slevc, 2019). This debate is accentuated when considering holistic music training, where musical and non-musical aspects intertwine, blurring near vs. far transfer boundaries. The distinction challenge originates from research implementation. Targeting specific music components (e.g., pitch, rhythm) clarifies mechanisms and transfer boundaries, while holistic training mimics natural learning but lacks this precision. Current research emphasizes holistic training, leaving gaps in understanding mechanisms for music-induced transfer. To enhance comprehension, investigating both holistic and component-based training offers insights for educational and health applications and technology development.

Our findings have practical implications for various domains: education, clinical populations, auditory perception, language/communication, social functioning, and emotion/reward. Inhibition control is pivotal for child development, encompassing self-regulation, reasoning, focus, and reducing compulsive behavior (Allom et al., 2016; Bowley et al., 2013; Fogel et al., 2019; Liu et al., 2015; Palermo & Bartoli, 2019; Wilkinson et al., 2020). Music training, being rewarding and motivating, could complement general education (Bogacz, 2020; Di Domenico & Ryan, 2017; Koops & Kuebel, 2018). Music might efficiently enhance inhibition control, especially in neurodevelopmental conditions, offering enjoyable contexts for learning (Takacs & Kassai, 2019). Autism interventions without a social component also show benefits (Diamond, 2013; Pasqualotto et al., 2021; Srinivasan et al., 2015, 2016). Music’s neural overlap with language influences language skills; inhibition training aids language switching (Wu et al., 2021). Rhythm training also improves language skills (Flaugnacco et al., 2015; Gordon et al., 2011, 2015) through neural entrainment models (Large & Jones, 1999; Nozaradan, 2014). Pitch skills link to phonological awareness (Loui et al., 2012; Patscheke et al., 2019), which is essential for reading. Music enhances socio-emotional development (Buren et al., 2021; Gaudette-Leblanc et al., 2021; Keller et al., 2014). Inhibition control’s role in emotions, turn-taking, and emotional abilities in turn influences social and emotional growth (Brodal et al., 2017; Nummenmaa et al., 2021; Reybrouck et al., 2018; Rose-Krasnor & Denham, 2009; van Noorden et al., 2017; Vuust et al., 2011). In sum, inhibition control mediates skill transfer from music training to non-musical activities, enhancing communication, impulse control, and socio-emotional skills. Music training complements education and clinical interventions effectively.

This meta-analysis shows a robust overall effect among longitudinal designs and is sufficiently powered to assert the ecological validity of our results. We found a moderating effect of the intervention environment for music training on inhibition control in children, indicating training in a music facility is more impactful than training at an elementary school. This effect is likely due to better-trained music instructors, a more positive environment conducive to music learning, and specialized music facilities and equipment. Our effect sizes remain significant and robust across ages, training intensity, and randomization. The absence of an age effect is likely due to the overall narrow age range and alignment with the sensitive period for music training (Penhune, 2011). Surprisingly, training intensity did not moderate skill transfer to inhibition control in children, possibly due to training focus and sufficient lower training time. The results imply that over 300 minutes of training in varied environments yield inhibition control improvements. Mixing community-based, quasi-experimental, and RCT designs adds ecological validity to our results and helps assess the generalizability of our conclusions (Habibi et al., 2022). The analysis of 8 studies using RCT designs in this meta-analysis yielded a greater effect size estimate than the analysis of all longitudinal studies. The fact that the RCT studies found the greatest improvement in inhibition control outcomes for music training emphasizes the importance of strong experimental design and supports future focus on RCT designs for studying music training. RCTs comprised 36% of the included studies and had a greater overall effect size than the superset of longitudinal studies.

Nonetheless, the set of studies is relatively small, and future RCTs will benefit the field and establish the causality of music training interventions on executive function outcomes with further certainty. Given the nature of the comparison between musical and non-musical interventions, only single blinding was possible. Future studies should also conduct follow-up testing to assess the temporal stability of inhibition control improvements after music training. A study by Bentley et al. (2023) shows that the beneficial effects on inhibition control in Williams et al. (2023) were stable and could be captured six months later, thus encouraging future work to conduct follow-up examinations.

Importantly, music training/interventions can yield varied effects across individuals. Assessing how additional factors like age, baseline performance, or clinical severity influence training outcomes is crucial. Designs that consider individual differences offer targeted, personalized training/intervention recommendations, such as adjusting the difficulty to optimize motivation and cognitive efficiency (Ryan & Deci, 2017; Bogacz, 2020). Computerized training allows flexibility for diverse users, including those with cognitive impairments. By combining individual differences and technology, adaptive training can overcome barriers and enhance potential (Karbach et al., 2017; Katz et al., 2016). Parkinson’s disease research from our laboratory underlines the significance of this approach (Dalla Bella et al., 2017, 2018; Puyjarinet et al., 2022). Embracing individual differences and tailored approaches holds promise for efficient and accessible music training in the future.

Further research incorporating these important design criteria will help to clarify the mechanisms and necessary conditions of this transfer effect. Additionally, a specific focus on the contribution of the core elements of music, such as pitch, rhythm, and non-social music training, will help determine what components of music training most drive this effect. Because inhibition control is a developmentally fundamental executive function and is important for language/communication abilities and socio-emotional skills, studying inhibition as a potential mediating factor for improving outcomes in education and clinical populations with socio-communicative deficits is a promising avenue of research. Using theory-based and mechanism-driven predictions for specific targeted outcomes will help develop the potential of music training to strengthen children’s cognitive abilities and complement remediation and teleremediation in clinical populations.

## Methods

### Inclusion criteria

In this meta-analysis, we included studies that met the following criteria:

1. Music training was in the human neurotypical childhood and adolescent population.
2. The studies included an experimental group trained using music alone. Music was an activity that involved a production aspect using a musical instrument such as singing or drumming, or the music training required perceptual/motor skills related to music, such as in a computerized music training program.
3. The studies involved training music abilities.
4. The studies were longitudinal, with either an active or passive control group comparison.
5. The studies included a performance-based inhibition control measure as an outcome.
6. The studies were in peer-reviewed journals from January 1980 to December 2023.

### Searching procedure and collection process

We conducted an exhaustive literature search for music training and inhibition control studies in children and adolescents in various databases such as PubMed, PsychInfo, Web of Science, and Scopus. The following search keywords to cover music training were included: “music* training or music* learning or music* education or music* lessons or musician*”; and search keywords to cover inhibition control included: “executive* or executive function* or executive control or cognitive function* or cognitive control or inhibition”; and search keywords to cover development included: “development* or child or learning or cognitive development or childhood or children or adolescent*.” The search results were exported into EndNote software (version X20; Clarivate Analytics, New York, USA). After duplicates were removed, 2344 studies remained. A search on Google identified three additional unpublished manuscripts making a total of 2347 screened records. A Preferred Reporting Items for Systematic Reviews and Meta-Analyses (PRISMA; Liberati et al., 2009) diagram summarizing the number of studies meeting the search criteria is shown in Figure 1. The first author XX, a doctoral candidate with a MSc, evaluated these studies based on title and abstract and found 629 studies relevant to music training. For this procedure, if the title and abstract did not provide sufficient information, XX decided based on the full content of the publication. XX then screened all musically relevant studies based on title and abstract and determined that 237 studies had longitudinal designs, 216 had cross-sectional designs and 176 records did not fall in either category. After screening all the longitudinal studies based on the title and abstract, XX retained 44 studies that used inhibition control measures in children for full-text review. Following this, XX and the second author (YY), a research associate with a PhD, evaluated the full content of 44 studies, and 22 were excluded (see Supplemental Table A). Independent coding was carried out by XX and YY on the included 22 studies for quality assessment, and the resulting tables were compared and discussed to reach a consensus. Of the 22 included studies, 8 involved participant-level randomizations with an active control group and were classified as RCTs.

### Tasks & Measures

The present work targeted studies with inhibition control measures involving prepotent response inhibition and resistance to distractor interference. We included the following inhibition control tasks: The Stop-Signal task, the Go/No-Go, the Eriksen Flanker, and the Stroop test. Similar adaptations of these tasks were included, for example in pre-schoolers (Bowmer et al., 2018; Brown et al., 2022; Kosokabe et al., 2021). Each of these tasks had different primary outcome variables for measuring inhibition. Some, like the Go/No-Go and the Stroop test, recorded accuracy scores and focused on the percentage of correct withheld responses. Paradigms like the Eriksen Flanker task typically use correct response times as indicators of cognitive resource allocation. For example, trials with more incongruent information show slower response times than trials with congruent information. When the means of these two trial types are subtracted from each other, the remaining response time represents the added cognitive delay of resolving conflicting information. The smaller this value is, the greater the participant’s inhibition control and resistance to distractor interference.

For this analysis, we calculated effect size estimates for accuracy and reaction times based on the most suitable measure for a meta-analysis. In the event of missing data, we contacted the corresponding authors of the respective studies three times. We subsequently omitted studies if authors did not respond, or if means and standard deviations were not reported or calculable. When studies only provided data visualizations of means and standard deviations, we manually extracted numerical data using WebPlotDigitalizer (http://automeris.io/WebPlotDigitizer), a tool designed to estimate values from graphical visualizations. In addition to collecting data on inhibition control measures, we coded the following information for each study:

a. Age
b. Music training intensity (number of sessions, session durations, total minutes of training)
c. Context of the training (school-based or a specialized music organization)
d. Participant-level randomization

Specific decisions about inhibition control measures for analysis are shown below in Table 1.

### Unpublished studies

There are arguments for and against including unpublished studies in meta-analytic methodology (Sterling et al., 1995). When sourcing studies only from peer-reviewed journals, a risk of publication bias increases in favor of studies with significant results, thus inflating the global effect size (Rosenthal & DiMatteo, 2001). In contrast, including studies that have not undergone a rigorous peer-reviewed process may decrease the overall quality of study designs and interpretability of the findings. Given the 42-year scope of the current meta-analysis, obtaining a representative sample of unpublished works was not feasible due to technical limitations (i.e., contact information no longer valid, research institutes or laboratories closed). We carried out search queries for this meta-analysis on Scopus and ProQuest Dissertations and found one relevant study. We decided not to include unpublished studies in the present work.

### Moderator variable coding

We coded and analyzed a range of potential moderators to assess if the variations in the effect of music training on inhibition control depended on these variables. The mean participant age in each study was estimated based on available information. Training intensity was estimated as the total number of minutes in the music intervention based on the information in the published material. When information about intervention duration was given in terms of school years, unless specified otherwise, we used 32 weeks per year to estimate the training intensity. Training environment was classified based on whether music training occurred at school, or in a context outside the school environment. Randomization was classified based on whether participants were individually randomized to a training group.

### Quality Assessment

To assess the potential impact of bias in the reviewed studies, we conducted a rigorous quality assessment based on the CONSORT criteria (Moher et al., 2010). For detailed information on individual studies, see Supplemental Table B, Supplemental Table C, Supplemental Table D, and Supplemental Table E. Specific information in this assessment included sample sizes, adherence to randomization, reported participant information (age, sex, IQ, socio-economic makeup, ethnic background), group matching status, attrition during training, training description, control group information, training intensity, outcome measure type, quality of music instruction, and participants’ prior music training. We assessed the quality of each study using the Cochrane Collaboration tool (Higgins et al., 2011). It is difficult to blind participants from knowing if they are in a music intervention, and no study in this analysis reached the quality level of a double-blind placebo-controlled design. As a result, all studies were of “moderate” quality based on the Cochrane standards.

### Attrition during training

Attrition was calculated separately by group in each study, based on the number of participants who started the training and who completed both the training and post-training measures. To assess potential attrition bias, differences in attrition between music and control groups in the full set of studies were evaluated using a mixed-effects model on these N values (i.e., interaction between timepoint and group).

### Quantitative meta-analysis

Following meta-analytic guidelines (Lipsey et al., 2001), we synthesized and analyzed the set of longitudinal studies. This procedure involved the following steps: describing relevant characteristics of studies in a table, calculating standardized mean difference effect sizes (Hedges’ g) for each comparison, determining overall effect size, and identifying potential moderator variables (Rosenthal, 1995; Rosenthal & DiMatteo, 2001). Hedges’ g effect sizes, reflecting the difference in pre-post performance change between groups in each study, were calculated using R version 4.2.0 software (R Core Team, 2022). In several studies, results were reported separately for participant age strata, or more than one performance measure was selected for inclusion in our meta-analysis. In these cases, we calculated individual effect sizes per age strata and/or measure, and then combined these to a single effect size for each study by running a supplemental fixed-effects meta-analysis using the “rma.mv “function and “FE” method of the “metafor” package 4.2.0 in R (Viechtbauer, 2010). The final meta-analyses of eight RCT studies and 20 longitudinal studies thus each contained one effect size per study. Effect sizes are described in the text as “small” (g ≅ 0.2), “medium/moderate” (g ≅ 0.5), or “large” (g ≅ 0.8) based on values suggested by Cohen (1988) and commonly used in studies of skill transfer effects in cognitive training (e.g., Bediou et al., 2018; Edwards et al., 2018; Sala et al., 2018).

Hedges’ g effect sizes were calculated using equation 8 of Morris (2008), who refer to this effect size as dppc2. This calculation incorporates Bessel’s correction as well as Hedges’ correction for small sample bias.

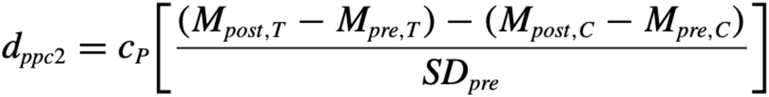

where

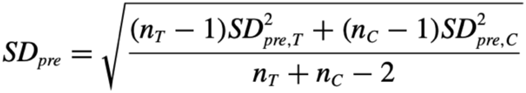

and

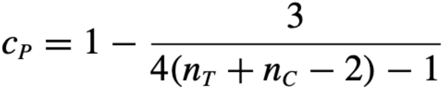

The variance of effect sizes was calculated using equation 25 of Morris (2008), where delta indicates the effect size calculated above:

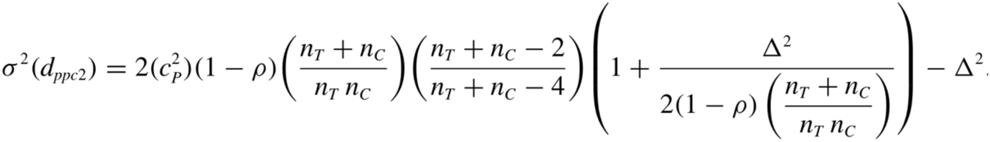

The variance equation requires specifying a correlation value (rho) between the pre-and post-training values. As this value was not reported by the published studies, we imputed a value of rho = .5 when performing the meta-analyses. We additionally verified that the significance of the meta-analysis results did not change if a value of rho = .001 or .999 is used instead.

Each meta-analysis was calculated in R using the “rma.mv” function in “metafor” (Viechtbauer, 2010). The “rma.mv” function performs a random-effects multilevel analysis that accounts for variation across studies and is recommended when studies vary in their samples or methodology. Moreover, random effects provide greater control for sample size differences when estimating effect sizes (Borenstein et al., 2009). We assessed between-study heterogeneity using the restricted maximum likelihood (REML) method, and confidence intervals of effect sizes were estimated using the Q-profile method, following the recommendations of (Veroniki et al., 2016).

Following analysis, we summarized heterogeneity across studies using the I^2^ measure (Higgins & Green, 2008). This measure provides an estimate of the proportion of variation of effect sizes that arises from clinical and methodological heterogeneity rather than chance and is recommended by Cochrane Reviews (Higgins et al., 2019). We created forest plots to display all the effect estimates and confidence intervals for individual studies and the combined meta-analytic result (Lewis & Clarke, 2001). These plots also provide a visual indication of heterogeneity among studies (Phan et al., 2015). We reported confidence intervals at the 95% level, p values were considered significant at values < 0.05, and we noted non-significant trends at p < 0.10, considering the low power of generally small sample sizes in the reviewed studies.

### Small study bias

Small study bias refers to the probability of small studies driving the overall effect size of the meta-analysis, thus impacting the validity of the findings. We used two techniques to determine the presence of a small study bias. First, as a visual diagnostic, funnel plots were generated to present the effect size of individual studies against the standard error associated with each study. Asymmetry around the triangular funnel provides an indication of potential publication bias (Rothstein et al., 2006). Second, we used Egger’s regression test (Egger et al., 1997) as a statistical evaluation of asymmetry, as implemented in the “regtest” function of the “metafor” package using a weighted regression model with a multiplicative dispersion term.

### Transparency and Openness

We adhered to the MARS guidelines for meta-analytic reporting (Appelbaum et al., 2018). The present work abides by the Transparency and Openness Promotion (TOP) at level 2 (Nosek et al., 2015):

1. Citation: Level 2, All data, program code, and other methods developed by others are appropriately cited in the text and listed in the references section.
2. Data Transparency: Level 2, processed data on which study conclusions are based are available and accessible on the anonymized OSF link: https://osf.io/d36gk/?view_only=8f57b9eef4fe42089f0a59ac1e03c26e
3. Analytic Methods (Code) Transparency: Level 2, computer code to reproduce analyses in an article is available on the anonymized OSF link: https://osf.io/d36gk/?view_only=8f57b9eef4fe42089f0a59ac1e03c26e
4. Research Materials Transparency: Level 2, Requirement—Materials described in the method section are available on the anonymized OSF link: https://osf.io/d36gk/?view_only=8f57b9eef4fe42089f0a59ac1e03c26e
5. Design and Analysis Transparency (Reporting Standards): Level 2, Authors complied with APA Style Journal Article Reporting Standards (JARS and MARS)
6. Citation: Level 2, All data, program code, and other methods developed by others are appropriately cited in the text and listed in the references section
7. Study Preregistration: Level 2, The study design and hypotheses for the present work were not preregistered. However, the study design and hypotheses, including half the study sample, were presented at the 16th Annual NeuroMusic Conference, 2020 and are publicly accessible at: [LINK DISCLOSES IDENTITY OF AUTHORS BUT IS IN COVER LETTER]
8. Analysis Plan Preregistration: Level 2, The study analysis plan for the present work was not preregistered. However, the analysis plan, including half the study sample, was presented at the 16th Annual NeuroMusic Conference, 2020 and are publicly accessible at: [LINK DISCLOSES IDENTITY OF AUTHORS BUT IS IN COVER LETTER]

## Notes

https://osf.io/d36gk/?view_only=8f57b9eef4fe42089f0a59ac1e03c26e

### Competing Interest Statement

The authors have declared no competing interest.

### Summary of Updates

Manuscript has been updated for 2024 and made more concise.

