## Supplemental Tables for "Does Music Training Improve Inhibition Control in Children? A Systematic Review and Meta-Analysis"

### Supplemental Materials

**Supplemental Table A**

*List of Excluded Studies and Reasons for Exclusion*

| Study | Reason | DOI / URL |
|---|---|---|
| McClelland et al. (2011) | Did not contain an active music condition | 10.1080/10409289.2011.574258 |
| Sportsman et al. (2012) | Did not have a performance IC measure | https://d.lib.msu.edu/etd/246 |
| Costa-Giomi (2015) | Did not have a performance IC measure | 10.1177/8755123314540661 |
| Schmitt et al. (2015) | Did not contain an active music condition | 10.1016/j.ecresq.2014.08.001 |
| Hedeyati et al. (2016) | Pre-training means and standard deviations were not reported and could not be obtained by contacting authors | 10.1080/13554794.2016.1241885 |
| Janus et al. (2016) | Did not have a performance IC measure | 10.1016/j.jecp.2015.11.009 |
| Sachs et al. (2017) | Study had the same cohort as Hennessy et al. (2019) but presented a smaller set of results | 10.1371/journal.pone.0187254 |
| Holochwost et al. (2017) | Control group data were not reported and could not be obtained by contacting authors | 10.1037/aca0000112 |
| Aleman et al. (2017) | No pre-post means reported – author contacted three times and did not respond | 10.1007/s11121-016-0727-3. |
| Maróti et al. (2019) | Did not compare music to a non-music activity | 10.1177/0305735618778765 |
| McClelland et al. (2019) | Did not contain an active music condition | 10.3389/fpsyg.2019.02365 |
| Norgaard et al. (2019) | Did not compare music to a non-music activity | 10.1177/0022429419863038 |
| Welch et al. (2021) | Thesis: Not published | https://www.proquest.com/openview/e9775ce195663e12068f834093ea379c/1?pq- |

**Supplemental Table A**

*List of Excluded Studies and Reasons for Exclusion*

| Study | Reason | DOI / URL |
|---|---|---|
| | | origsite=gscholar&cbl=44156 |
| Bayanova et al. (2022) | Pre- and post-training means and standard deviations were not reported and could not be obtained by contacting authors | 10.3390/educsci12020119? |
| Hogan et al. (2018) | Did not compare music to a non-music activity | 10.1016/j.ecresq.2017.12.004 |
| Kasuya-Ueba et al. (2020) | Did not have a performance IC measure | 10.3389/fnins.2020.00757 |
| Remache et al. (2022) | Spanish thesis: Not published | https://orcid.org/0000-0001-7771-1036 |
| Bentley et at. (2023) | Follow-up study to Williams et al. (2022) | 10.1111/desc.13358 |
| Hogan et al. (2023) | Did not compare music to a non-music activity | 10.1037/aca0000593 |
| Schibli et al. (2023) | Pre-training means and standard deviations were not reported and could not be obtained by contacting authors | 10.3390/brainsci13050783 |
| Barnard et al. (2023) | Thesis: Not published | https://www.proquest.com/openview/f553d65a35ab9b4dc28100598cb3c820/1?pq-origsite=gscholar&cbl=18750&diss=y |
| Zachariou et al. (2023) | Did not compare music to a non-music activity | 10.1007/s11409-023-09342-1 |

**Supplemental Table B**

*Qualitative Analysis of Studies (Intervention Details)*

| First author | Year | Music condition | Led by music teacher | Control condition | Prior music training |
|---|---|---|---|---|---|
| Bolduc | 2020 | Rhythmic-centered training | ✓ | Free and guided play with motor movement | N.R. |
| Bowmer | 2018 | Singing, body synchronization | ✓ | Visual arts | N.R. |
| Brown | 2022 | Multi-component integrated music training | ✓ | Usual preschool | 45% of music group had received prior training; does not mention prior music training of control group |
| Bugos | 2017 | Multi-component integrated music training | ✓ | Lego training | No prior music training |
| Dégé | 2020 | Multi-component integrated music training | Instructors received training from a manual | Sports | N.R. |
| D'Souza | 2018 | Multi-component integrated music training | ✓ | Dance training | N.R. |
| Fasano | 2019 | Multi-component integrated music training | ✓ | Passive | No prior music training |
| Frischen | 2019 | Two separate pitch and rhythm training combined | Instructors received training from a manual | Sports | 16% had prior early-music education; matched between |
| Frischen | 2021 | Multi-component integrated music training | Instructors received training from a manual | Visual arts | No prior music training |
| Guo | 2018 | Pitch-based | ✓ | Usual play break | Excluded for current music lessons |
| Hallberg | 2017 | Violin Suzuky Method | ✓ | Waitlist | N.R. |

**Supplemental Table B**

*Qualitative Analysis of Studies (Intervention Details)*

| First author | Year | Music condition | Led by music teacher | Control condition | Prior music training |
|---|---|---|---|---|---|
| Hennessy | 2019 | Multi-component integrated music training | ✓ | Sports | Current music/sports exclusion above a certain intensity |
| Jaschke | 2018 | Multi-component integrated music training | ✓ | Visual arts | No prior music training |
| Kosokabe | 2022 | Rhythmic-centered training | × | Drama training | N.R. |
| Linnavalli | 2018 | Multi-component integrated music training | ✓ | Dance training | Some participants had prior music training |
| Moreno | 2011 | Perceptual/motor music-based computer training | ✓ | Visual arts | N.R. |
| Shen | 2019 | Perceptual/motor music-based computer training | ✓ | Passive | No prior music training |
| Sperling | 2023 | Multi-component integrated music training | N.R. | Passive | N.R. |
| Suppalarkbunlue | 2022 | Rhythmic-centered training | Instructors received training from a manual | Usual preschool | No prior music training at preschool; private lessons N.R. |
| Vazou | 2020 | Rhythmic-centered training | Instructors received training from a manual | Sports | N.R. |
| Williams | 2019 | Rhythmic-centered training | ✓ | Usual preschool | N.R. |
| Williams | 2023 | Rhythmic-centered training | × | Usual preschool | N.R. |

**Supplemental Table C**

*Qualitative Analysis of Studies (Matching and Demographics)*

| First author | Year | Music N ≥ 20 | Control N ≥ 20 | IQ matched | Sex matched | SES matched | Age matched | % Female reported |
|---|---|---|---|---|---|---|---|---|
| Bolduc | 2020 | ✓ | ✓ | ✓ | ✓ | ✓ | ✓ | 53 |
| Bowmer | 2018 | × | × | ✓ | × | N.R. | ✓ | 72 |
| Brown | 2022 | ✓ | ✓ | N.R. | × | ✓ | ✓ | 59 |
| Bugos | 2017 | × | × | N.R. | ✓ | ✓ | ✓ | 56 |
| Dégé | 2020 | × | × | ✓ | ✓ | ✓ | ✓ | 60 |
| D'Souza | 2018 | ✓ | ✓ | ✓ | ✓ | ✓ | ✓ | 59 |
| Fasano | 2019 | ✓ | ✓ | N.R. | ✓ | ~ | ✓ | 50 |
| Frischen | 2019 | ✓ | ✓ | ✓ | ✓ | ✓ | ✓ | 61 |
| Frischen | 2021 | ✓ | ✓ | ✓ | ✓ | ✓ | ✓ | 52 |
| Guo | 2018 | ✓ | ✓ | ✓ | ✓ | N.R. | ✓ | 45 |
| Hallberg | 2017 | ✓ | ✓ | N.R. | N.R. | N.R. | ✓ | N/R |
| Hennessy | 2019 | × | × | ✓ | ✓ | ✓ | ✓ | 41 |
| Jaschke | 2018 | ✓ | ✓ | ✓ | ✓ | ✓ | ✓ | 52 |
| Kosokabe | 2022 | ✓ | ✓ | N.R. | ✓ | N.R. | ✓ | 55 |
| Linnavalli | 2018 | × | × | N.R. | ✓ | ✓ | ✓ | 54 |
| Moreno | 2011 | ✓ | ✓ | ✓ | ✓ | ✓ | ✓ | 59 |
| Shen | 2019 | ✓ | ✓ | ✓ | ✓ | N.R. | ✓ | 61 |
| Sperling | 2023 | ✓ | ✓ | N.R. | ✓ | ✓ | ✓ | 55 |

**Supplemental Table C**

*Qualitative Analysis of Studies (Matching and Demographics)*

| First author | Year | Music N ≥ 20 | Control N ≥ 20 | IQ matched | Sex matched | SES matched | Age matched | % Female reported |
|---|---|---|---|---|---|---|---|---|
| Suppalarkbunlue | 2022 | ✓ | ✓ | N.R. | ✓ | N.R. | ✓ | 52 |
| Vazou | 2020 | ✓ | × | ✓ | ✓ | N.R. | ✓ | 46 |
| Williams | 2019 | ✓ | ✓ | N.R. | ✓ | ✓ | ✓ | 48 |
| Williams | 2023 | ✓ | ✓ | N.R. | ✓ | ✓ | ✓ | 51 |

* ~ = marginal p 0.05 ~ 0.10

**Supplemental Table D**

*Qualitative Analysis of Studies (Socio-Economic and Cultural Makeup, and Attrition)*

| First author | Year | Socio-economic makeup | Cultural/ethnic makeup | Country | Attrition (%) Music group | Attrition (%) Control group |
|---|---|---|---|---|---|---|
| Bolduc | 2020 | Middle-class | N.R. | Canada (French) | 14 | 10 |
| Bowmer | 2018 | 24.4% low income based on the meal plan | 43.9% were not fluent in English living in West London | England | 0 | 0 |
| Brown | 2022 | Mean family income was 20 000 USD per year | 18 % were Caucasian | United States of America | 0 | 0 |
| Bugos | 2017 | Income range 10 000$-120 000$; 57% of parents had no more than a high-school degree | 37.5% Caucasian 57% reporting | United States of America | 6 | 6 |
| Dégé | 2020 | At least 78% of kindergarten parents had a university degree in this study | N.R. | Germany | 0 | 0 |
| D'Souza | 2018 | All mothers had a college degree | All monolinguals in Canada | Canada | 0 | 0 |
| Fasano | 2019 | At least 11% of both parents unemployed | N.R. | Italy | 0 | 0 |
| Frischen1 | 2019 | At least 18% of study sample below minimum wage for family income | N.R. | Germany | 20 | 23 |
| Frischen2 | 2021 | ~10% of study sample below minimum wage for family income | N.R. | Germany | 25 | 14 |
| Guo | 2018 | All children attended a | N.R. | Japan | 0 | 0 |

**Supplemental Table D**

*Qualitative Analysis of Studies (Socio-Economic and Cultural Makeup, and Attrition)*

| First author | Year | Socio-economic makeup | Cultural/ethnic makeup | Country | Attrition (%) Music group | Attrition (%) Control group |
|---|---|---|---|---|---|---|
| | | public school; no mention of income range | | | | |
| Hallberg | 2017 | 46 % of school students below national poverty line | Excluded children from a non-English home environment | United States of America | 0 | 0 |
| Hennessy | 2019 | All participants from underprivileged communities in Los Angeles | 97.7% of participants were bilingual and spoke either Spanish or Korean in addition to English | United States of America | 0 | 0 |
| Jaschke | 2018 | No mention of family income; All parents had at least a high-school diploma | N.R. | Netherlands | 5 | 34 |
| Kosokabe | 2022 | Middle-class | N.R. | Japan | 0 | 0 |
| Linnavalli | 2018 | No mention of family income; All mothers had at least a high-school diploma | N.R. | Finland | 0 | 0 |
| Moreno | 2011 | Mother's average education was a bachelor's college degree | N.R. | Canada | 26 | 26 |
| Shen | 2019 | Middle-class | N.R. | China | 0 | 0 |
| Sperling | 2023 | Participants mostly from disadvantaged communities | 44% Latin-American, 40% African-American, 11% Caucasian, 7% Other | United States of America | N.R.* | N.R.* |

**Supplemental Table D**

*Qualitative Analysis of Studies (Socio-Economic and Cultural Makeup, and Attrition)*

| First author | Year | Socio-economic makeup | Cultural/ethnic makeup | Country | Attrition (%) Music group | Attrition (%) Control group |
|---|---|---|---|---|---|---|
| Suppalarkbunlue | 2022 | N.R. | N.R. | Thailand | N.R.* | N.R.* |
| Vazou | 2020 | N.R. | 77% Caucasian | United States of America | 8 | 0 |
| Williams | 2019 | Participants mostly from disadvantaged communities | At least 17.4% of sample had an Aboriginal, Torres Islander, or non-English speaking background | Australia | N.R.* | N.R.* |
| Williams | 2023 | Participants mostly from disadvantaged communities | At least 17.4% of sample had an Aboriginal, Torres Islander, or non-English speaking background | Australia | 0 | 2 |

*Reported attrition information did not permit calculating per-group values in the context of this meta-analysis

**Supplemental Table E**

*Descriptive Summary of Studies Included in the Meta-Analysis*

| First author | Year | Mean age [a] | Age range [a] | Selected measure(s) | Total training (min) | Music (N) [b] | Control (N) [b] | Participant-level randomized group assignment |
|---|---|---|---|---|---|---|---|---|
| Bolduc | 2020 | 5.5 | 4-6 | NEPSY-II inhibition | 760 | 50 | 102 | ✓ |
| Bowmer | 2018 | 3.8 | 3-4 | Peg Tapping | 320 | 15 | 27 | ✓ |
| Brown | 2022 | 4.0 | 3-5 | Combined Z score for 3 inhibition tasks for preschoolers | 720 | 148 | 191 | × |
| Bugos | 2017 | 4.9 | 4-5 | Night Stroop | 540 | 17 | 34 | ✓ |
| Dégé | 2020 | 6.0 | | NEPSY-II inhibition | 840 | 11 | 25 | ✓ |
| D'Souza | 2018 | 7.6 | 6-9 | Stroop | 1800 | 24 | 50 | ✓ |
| Fasano | 2019 | 8.9 | 8-10 | Stop-Signal Walk-No-Walk | 1200 | 55 | 113 | × |
| Frischen | 2019 | 5.7 | 5-6 | NEPSY-II inhibition | 1200 | 53 | 76 | ✓ |
| Frischen | 2021 | 6.6 | 6-7 | NEPSY-II inhibition | 1440 | 27 | 58 | ✓ |
| Guo | 2018 | 7.5 | 6-8 | Go/No-Go | 300 | 20 | 40 | ✓ |
| Hallberg | 2017 | 5.2 | N.R. | KCCP Test 5 - Commission errors | 900 | 26 | 48 | ✓ |
| Hennessy | 2019 | 6.8 | 6-7 | Flanker | 19500 | 17 | 32 | × |
| Jaschke | 2018 | 6.4 | 6-8 | Go/No-Go | 6480 | 42 | 71 | × |
| Kosokabe | 2022 | 4.7 | 3-5 | Combined Z scores (2 preschool inhibition tasks) | 900 | 92 | 143 | × |
| Linnavalli | 2018 | 5.3 | 5-6 | NEPSY-II inhibition | 2700 | 15 | 63 | × |
| Moreno | 2011 | 5.3 | 4-6 | Go/No-Go | 1800 | 24 | 48 | ✓ |
| Shen | 2019 | 4.2 | 4-5 | Day/Night Stroop | 2100 | 30 | 61 | × |
| Sperling | 2023 | 5.4 | 5-6 | Go/No-Go | 21600 | 49 | 60 | |

**Supplemental Table E**

*Descriptive Summary of Studies Included in the Meta-Analysis*

| First author | Year | Mean age [a] | Age range [a] | Selected measure(s) | Total training (min) | Music (N) [b] | Control (N) [b] | Participant-level randomized group assignment |
|---|---|---|---|---|---|---|---|---|
| Suppalarkbunlue | 2022 | 4.4 | 4-5 | Go/No-Go | 1080 | 39 | 79 | × |
| Vazou | 2020 | 7.7 | 6-11 | Flanker | 420 | 22 | 39 | × |
| Williams | 2019 | 4.7 | 4-5 | Go/No-Go | 480 | 59 | 113 | × |
| Williams | 2023 | 4.2 | 4-5 | Go/No-Go | 360 | 112 | 102 | |

[a] Years

[b] Analyzed

**Computer Code**

Runnable computer code can be found at the following URL:

https://osf.io/d36gk/?view_only=8f57b9eef4fe42089f0a59ac1e03c26e
